# Simultaneous stereo-EEG and high-density scalp EEG recordings to study the effects of intracerebral stimulation parameters

**DOI:** 10.1101/2021.11.15.468625

**Authors:** S. Parmigiani, E. P. Mikulan, S. Russo, S. Sarasso, F. M. Zauli, A. Rubino, A. Cattani, M. Fecchio, D. Giampiccolo, J. Lanzone, P. D’Orio, M. del Vecchio, P. Avanzini, L. Nobili, I. Sartori, M. Massimini, A. Pigorini

**Affiliations:** Department of Biomedical and Clinical Sciences “L. Sacco” Università degli Studi di Milano, Milan, Italy; Department of Philosophy “Piero Martinetti”, Università degli Studi di Milano, Milan, Italy; “C. Munari” Epilepsy Surgery Centre, Department of Neuroscience, Niguarda Hospital, Milan, Italy; Department of Mathematics & Statistics, Boston University, Boston, MA, USA; Center for Neurotechnology and Neurorecovery, Department of Neurology, Massachusetts General Hospital, Boston, MA, USA; Section of Neurosurgery, Department of Neuroscience, Biomedicine and Movement Sciences, University of Verona, Verona, Italy; Deparment of Systems Medicine, Neuroscience, University of Rome Tor Vergata, Rome, Italy; Istituti Clinici Scientifici Maugeri, IRCCS, Neurorehabilitation Department of Milano Institute, Milan, Italy; Istituto di Neuroscienze, Consiglio Nazionale delle Ricerche, Parma, Italy; Child Neuropsychiatry, IRCCS Istituto G. Gaslini, Genova, Italy; Istituto Di Ricovero e Cura a Carattere Scientifico, Fondazione Don Carlo Gnocchi, Milan, Italy; Azrieli Program in Brain, Mind and Consciousness, Canadian Institute for Advanced Research, Toronto, Canada

**Keywords:** Single pulse electrical stimulation, stereo-EEG, scalp hd-EEG, CCEP, stimulation parameters

## Abstract

**Background:** Cortico-cortical evoked potentials (CCEPs) recorded by stereo-electroencephalography (SEEG) are a valuable clinical tool to investigate brain reactivity and effective connectivity. However, these invasive recordings are spatially sparse since they depend on clinical needs. This sparsity hampers systematic comparisons across-subjects, the detection of the whole-brain spatiotemporal properties of CCEPs, as well as their relationships with classic sensory evoked potentials.

**Objective:** To demonstrate that CCEPs recorded by high-density electroencephalography (hd-EEG) are sensitive to changes in stimulation parameters and compensate for the limitations typical of invasive recordings.

**Methods:** SEEG and hd-EEG activities were simultaneously recorded during SPES in drug-resistant epileptic patients (N=36). Changes in stimulation parameters encompassed physical (pulse intensity and width), geometrical (angle and position with respect to white/grey matter) and topological (stimulated cortical area) properties. Differences were assessed by measuring the overall responses and the amplitude of N1 and N2 components of the CCEPs, and by their spectral profiles.

**Results:** While invasive and non-invasive CCEPs were generally correlated, differences in pulse duration, angle and stimulated cortical area were better captured by hd-EEG. Further, hd-EEG responses to SPES reproduced basic features of responses to transcranial magnetic stimulation and showed a much larger amplitude as compared to typical sensory evoked potentials.

**Conclusions:** The present results show that macroscale hd-EEG recordings are exquisitely sensitive to variations in SPES parameters, including local changes in physical and geometrical stimulus properties, while providing valuable information about whole-brain dynamics. Moreover, the common reference space across subjects represented by hd-EEG may facilitate the construction of a perturbational atlas of effective connectivity.

**Highlights:**

- CCEPs recorded with hd-EEG and SEEG are correlated.
- hd-EEG recording is highly sensitive to changes in stimulation parameters.
- hd-EEG responses show higher amplitude responses with respect to non-invasive ones.
- Simultaneous recordings provide a fixed observation point across subjects.

## Introduction

Intracortical electrical stimulation is an invaluable tool for surgical planning [1–3] and provides a direct assessment of brain evoked reactivity and effective connectivity in humans [4–6]. Clinical protocols often combine Single Pulse Electrical Stimulation (SPES) with stereotactic electroencephalography (SEEG) to evoke responses in areas explored with intracerebral electrodes [7,8]. Conceived for localizing the origin and diffusion of the epileptogenic activity [9–12] in patients with focal drug-resistant epilepsy, SPES typically elicits consistent cortico-cortical evoked potentials (CCEPs) whose features reflect physiological and pathological characteristics of the underlying neural tissue [7–9,13,14].

Thanks to their high functional specificity [15], signal fidelity [16], and high spatial and temporal resolution [12–14], CCEPs can be used as an electrophysiological tool to assess brain reactivity and effective connectivity complementing functional and structural connectivity measures [4,13,17,18]. However, invasive recordings are necessarily sparse since intracerebral electrodes are typically circumscribed to a limited set of brain regions differing from one subject to another depending on clinical needs [8,9,12,14,19]. The variability and sparsity of electrode placement clearly restricts a systematic comparison across subjects, the detection of the stimulation effects at the whole-brain level, as well as a direct comparison between CCEPs and other EEG potentials such as those evoked by non-invasive sensory, electrical or magnetic stimulation.

In the present work, we overcame these limitations by simultaneously acquiring high-density EEG recordings, which provide a fixed observation point to reliably compare the responses evoked by SPES across subjects and to assess their whole-brain dynamics, as well as their amplitude at the scalp level. Specifically, we analyzed the effects induced by the systematic manipulation of different stimulation parameters on CCEPs recorded from both SEEG and scalp EEG during wakefulness. We assessed CCEPs changes associated to physical (pulse intensity and width), geometrical (angle and position with respect to white/gray matter) and topological (stimulated cortical area) stimulation properties and found that, when compared to SEEG, high-density scalp EEG detects specific patterns that are more consistent across subjects. Notably, the differences in the overall response when stimulating different topological areas were systematically captured only by scalp recordings. We also observed a rostro-caudal gradient of the spectral properties of CCEPs evoked by the stimulation of different cortical areas, confirming previous results with Transcranial Magnetic Stimulation combined with EEG (TMS-EEG) studies [20]. Further, comparing the absolute amplitude of clinical SPES-evoked EEG responses to the typical amplitude of somatosensory, visual, auditory, or TMS-evoked EEG potentials revealed that the former are the largest electrical responses that can be elicited in the awake human brain.

## Materials and Methods

### Participants

A total of 36 patients (median age=33±8 years, 21 female, Table S1) from the “Claudio Munari ‘‘ Epilepsy Surgery Center of Milan in Italy were enrolled in the study. All subjects had a history of drug-resistant, focal epilepsy, and were candidates for surgical removal/ablation of the seizure onset zone (SOZ). 31 patients did no showed any anatomical malformation in the MRI, while the other 5 patients showed small anatomical alterations (see Table S1). All patients had no neurological or neuropsychological deficits. The investigated hemisphere/s and electrodes location was decided based on electroclinical data and reported - for each subject - in Figure S1. All patients provided their Informed Consent in accordance with the local Ethical Committee (ID 348-24.06.2020, Milano AREA C Niguarda Hospital, Milan, Italy) and with the Declaration of Helsinki.

### Electrodes placement and localization

Electrodes placement was performed as reported in [8] while electrode localization and anatomical labelling was performed as in [21]. Detailed descriptions can be found in Supplementary Materials.

### Simultaneous SEEG and hd-EEG Recordings

During the 1-3 weeks of hospitalization, SEEG activity was continuously recorded through a 192-channel recording system (NIHON-KOHDEN NEUROFAX-1200) with a sampling rate of 1000Hz. All acquisitions were referenced to two adjacent contacts located entirely in white matter [22]. During their last day of hospitalization all subjects included in the present study underwent simultaneous scalp non-invasive recordings by means of high-density Electroencephalogram (hd-EEG - 256 channels, Geodesic Sensor Net, HydroCel CleanLeads). Placement of the hd-EEG net on the head was performed by trained neurosurgeons using sterile technique, following a precise step-by-step protocol: (1) sterilization of the net, (2) removal of the protective bandage from the subject’s head, (3) skin disinfection with Betadine and Clorexan, (4) positioning of the hd-EEG net, and (5) reduction of the impedances below 25-50 kOhm using conductive gel. An example of this setup is shown in Figure 1. Hd-EEG was then recorded at 1000 Hz sampling rate using an EGI NA-400 amplifier (Electrical Geodesics, Inc; Oregon, USA) referenced to Cz. SEEG and hd-EEG recordings were aligned using a digital trigger signal generated by an external trigger box (EMS s.r.l., Bologna, Italy). At the end of the simultaneous data acquisition, the spatial locations of hd-EEG contacts and anatomical fiducials were digitized with a SofTaxicOptic system (EMS s.r.l., Bologna, Italy) and coregistered with a pre-implant 3D-T1MRI. The net was then removed, and the skin was disinfected again.

**Figure 1.**
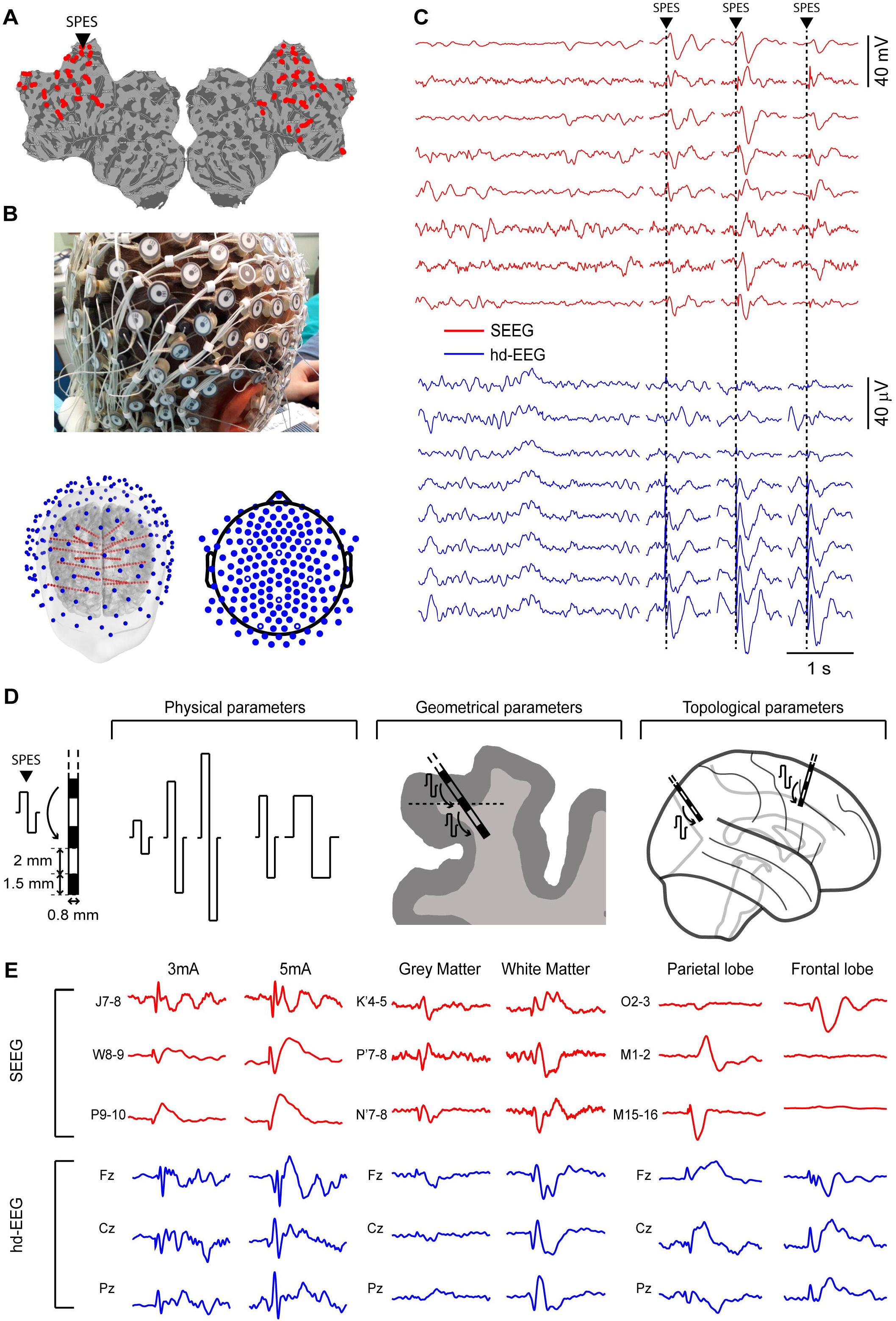
Experimental setup. **Panel A.** Topographical representation, on a flatmap, of the SEEG contacts in one representative subject. The black triangle indicates the contact used for SPES (anterior cingulate). **Panel B**. Picture of simultaneous SEEG and hd-EEG recordings, 3D reconstruction of the brain and SEEG implant of one representative patient (same as A) and topographical representation of hd-EEG contacts. **Panel C**. Concurrent raw intracerebral SEEG (red) and hd-EEG (blue) signals recorded respectively from eight representative bipolar contacts and from eight scalp EEG contacts. The black triangle and dashed vertical line indicate the time at which SPES was delivered. **Panel D.** Left, outline of a multi-lead intracerebral electrode. Right, overview of stimulation parameters categories: physical, geometrical and topological. **Panel E.** Examples of intracerebral SEEG (red) and hd-EEG (blue) signals recorded from representative bipolar contacts when delivering SPES with different physical (3mA vs 5mA), geometrical (white matter vs grey matter) and topological (parietal vs frontal lobe) parameters.

**Figure 2:**
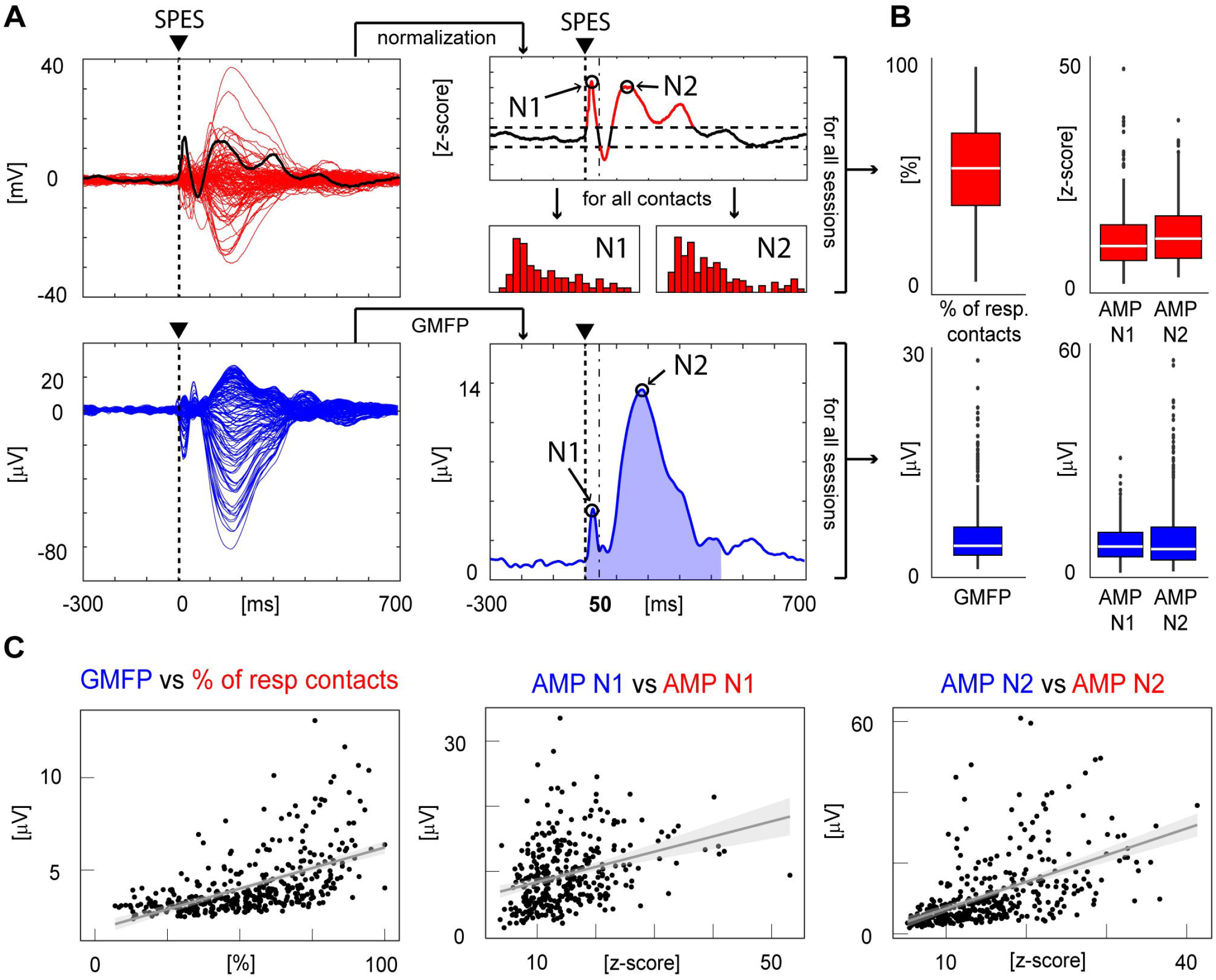
Quantification procedures and comparisons of hd-EEG and SEEG responses to SPES. **Panel A.** Butterfly plots, N1 and N2 detections, GMFP calculations and quantifications of SEEG (top panel, traces in red) and hd-EEG (bottom panel, traces in blue) responses to SPES. *SEEG:* the same procedure was performed for each significant SEEG bipolar contact (i.e. CCEP>6 STD of the baseline, as in [26]). After normalization (z-score) for the baseline (from −300ms to −50ms) and components detection (as in [35]), the amplitude of N1 and N2 components (black circles) were measured, obtaining two distributions (for N1 and N2). Values were then averaged across contacts to obtain average N1 and N2 amplitude values. *hd-EEG:* the GMFP is calculated from all hd-EEG contacts and then averaged between 0ms and 500ms (shaded blue area). The amplitude of N1 and N2 is detected as the maximum peak of the GMFP (black circles), respectively in the 0-50ms (dash and dot vertical line) and 50ms-300ms time window. **Panel B.** In red, from left to right, percentage of responding contacts and N1 and N2 amplitudes for all sessions, recorded in SEEG. In blue, from left to right, GMFP voltage and N1 and N2 amplitudes, for all sessions, recorded in hd-EEG. **Panel C.** Linear regression analyses comparing hd-EEG (on y-axes) and SEEG measures (on x-axes). From left to right, linear regression between GMFP calculated at the hd-EEG level and the number of SEEG contacts responding to SPES with a significant CCEP (*r*=0.592, *p*<0.001); linear regression between the amplitude of N1 component calculated both at the hd-EEG and at the SEEG level (*r*=0.313, *p*<0.001); linear regression between the amplitude of N2 component calculated both at the hd-EEG and at the SEEG level (*r*=0.553, *p*<0.001).

### Single Pulse Electrical Stimulation

During simultaneous hd-EEG and SEEG recordings, electrical single biphasic pulses (positive-negative) were injected between pairs of adjacent intracranial contacts pertaining to the same electrode with an inter-stimulus interval of at least one second across a wide range of intensities and pulse widths (see next paragraph). Brain activity was continuously recorded both from all other SEEG contacts as well as from the 256 scalp hd-EEG contacts. A single stimulation session consisted of 30/40 consecutive trials. The number of sessions varied between subjects (9 ± 4). All the sessions included in the present work (N=379) were selected following these criteria: stimulations (i) were delivered through a bipolar contact far from the SOZ (as indicated by electrical pathological activity and *a posteriori* confirmed by post-surgical assessment); (ii) were delivered through a bipolar contact that did not show spontaneous interictal epileptic activity (by visual inspection by P.dO., J.L., I.S.); (iii) did not elicit muscle twitches, somatosensory, or cognitive manifestations.

### Physical, geometrical, and topological stimulation parameters

This work includes a dataset collected in the context of presurgical evaluation during which SPES was delivered based on clinical assessment, thus employing different stimulation parameters. Retrospectively, we decided to group these parameters into three categories, namely *physical, geometrical* and *topological*.

*Physical stimulation parameters* included (i) stimulation intensity and (ii) pulse width. Stimulation intensities ranged from 0.1mA to 5mA. Specifically, SPES was delivered at 0.1mA (N=3), 0.5mA (N=13), 1mA (N=23), 3mA (N= 63) and 5mA (N= 223). Given the low number of sessions performed with intensities ≤1mA we decided to group together all these intensities (N=39). Pulse widths were instead 0.5ms (n=184) or 1ms (n=139).

*Geometrical stimulation parameters* refer (i) to the position of the stimulating bipolar contact with respect to the interface between grey matter and white matter and (ii) to the angle of insertion of the SEEG electrode with respect to the cortical surface. To derive both parameters we used the 3D meshes of the grey and white matter obtained with Freesurfer [25]. The distance to the grey/white matter boundary was computed as the distance between the center of the stimulating bipolar contact and the closest point on the white matter mesh (see Figure 1D and Figure 3A) using the *trimesh* library. The distances of the bipolar contacts were then lumped into three categories: both contacts in grey matter (*GG*), both contacts in white matter (*WW*) and one contact in grey matter and one in white matter (*GW*). The angle with respect to the cortical surface was calculated using the vector formed by the SEEG bipolar contact, and the normal vector of the closest segment of the white matter mesh (see Figure 1D and Figure 3C). Also the angles were lumped into two categories with respect to the cortical surface: *parallel* (δ < 45°; δ > 315°; and 135° < δ < 225°) and *perpendicular* (45° < δ < 135° and 225° < δ < 315°).

**Figure 3.**
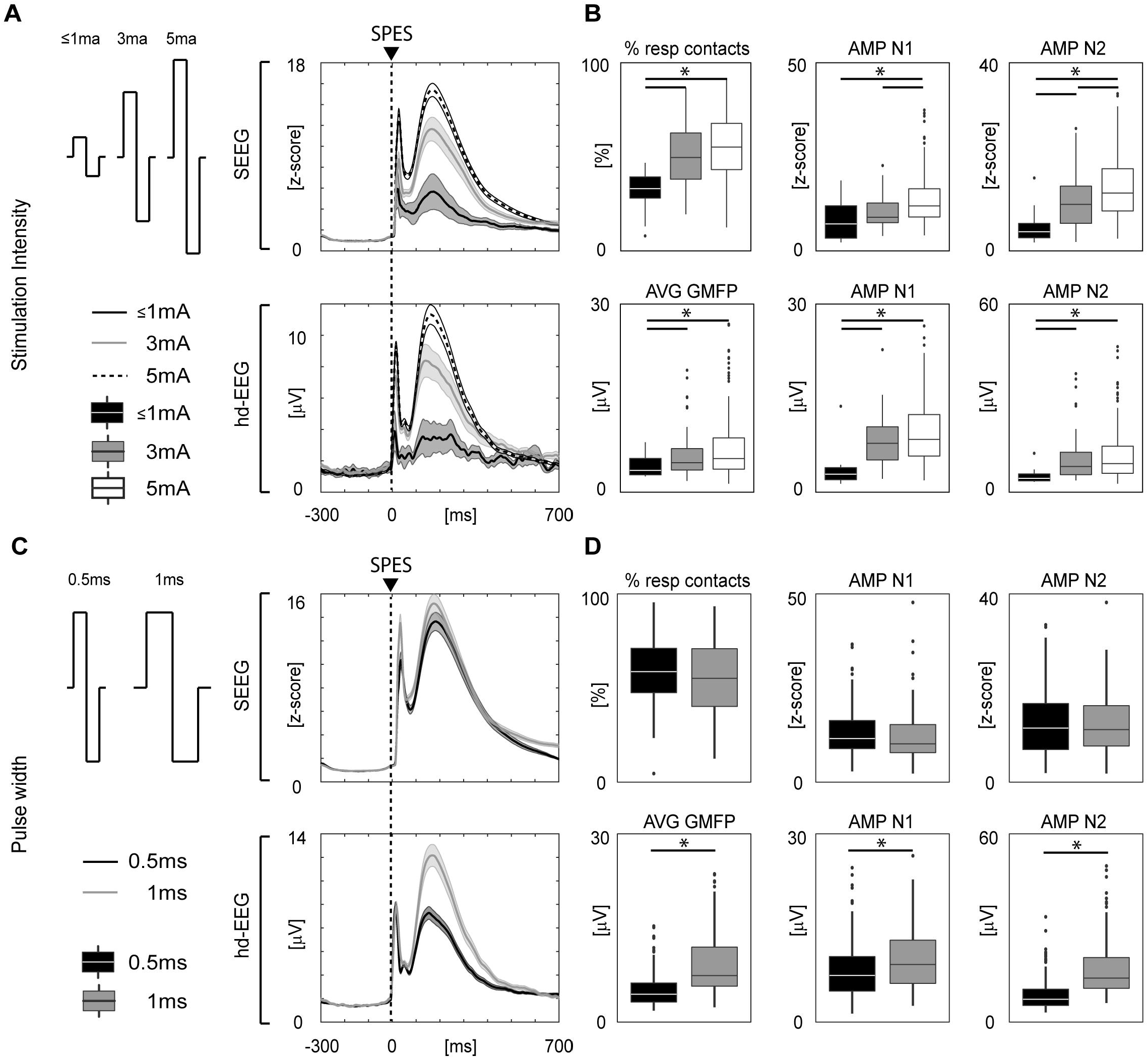
hd-EEG and SEEG responses to SPES delivered at different physical stimulation parameters: intensities and pulse durations. **Panel A.** Left, outline of the different pulse intensities; right, grand average of data obtained from all the subjects and sessions. **Panel B**. *Upper line:* from left to right, percentage of responding contacts and N1 and N2 amplitudes for all sessions, recorded in SEEG. *Lower line:* from left to right, GMFP voltage and N1 and N2 amplitudes, for all sessions, recorded in hd-EEG. Asterisks indicate significant statistical differences (post-hoc, two-tailed WR, p<0.01, FDR corrected). **Panel C**. Left, outline of the different pulse durations; right, grand average of data obtained from all the subjects and sessions. **Panel D.** *Upper line:* from left to right, percentage of responding contacts and N1 and N2 amplitudes for all sessions, recorded in SEEG. *Lower line:* from left to right, GMFP voltage and N1 and N2 amplitudes, for all sessions, recorded in hd-EEG. Asterisks indicate significant statistical differences (WR, p<0.01, FDR corrected).

*Topological stimulation parameters* included the following stimulated cortical areas: Cingulate cortex (n=30), Frontal cortex (n=93), Insula (n=26), Occipital cortex (n=37), Parietal cortex (n=113), Temporal cortex (n=80) using the Desikan-Killiany atlas for the anatomical labelling.

### Data pre-processing

The joint visual inspection of both SEEG and hd-EEG CCEPs allowed to retain 323/379 sessions (~85%), excluding 9 sessions that showed evoked epileptic spikes either at the scalp or intracerebral EEG level (Figure S1, see also [12]), and 47 sessions characterized by a number of retained trials lower than 25 due to overall bad data quality or the presence of interictal activity.

For the retained SPES sessions SEEG data were processed as in [24] while hd-EEG data were preprocessed as in [25]. Detailed procedures are reported in Supplementary Material.

### Amplitude Analysis

The effects of SPES parameters were assessed both at the SEEG and the hd-EEG level by measuring standard features of CCEP waveforms (amplitude of N1 and N2) as well as surrogate measures of the overall response (Figure 2). Specifically, at the SEEG level, the latter was quantified as the number of contacts responding with a significant CCEP (above 6 STD of the baseline, Figure 2B) to SPES [26], while the amplitudes of N1 and N2 were obtained at the single contact level [27,38] and then averaged across contacts. Conversely, at the hd-EEG level, the overall response was obtained as the Global Mean Field Power (GMFP, between 0ms and 500ms, shaded blue area in Figure 2B) and the amplitude of N1 and N2 were detected as the maximum peak of the GMFP (black circles in Figure 2B), respectively in the 0-50ms (dash and dot vertical line in Figure 2) and 50ms-300ms time window. SNR was calculated in the same time windows.

### Spectral Analysis

We performed an analysis similar to [20], in which we compared the spectral properties of the CCEPs elicited by the stimulation of the occipital, parietal and frontal cortices. Specifically, for each session, we conducted a time-frequency spectral analysis (Event Related Spectral Perturbations, ERSP [29]) that was averaged across contacts within each session. The resulting time-frequency power distribution was cumulated over time between 10ms and 150ms. Finally, due to the prominent presence of a non-specific response (*i.e*., the N1 - N2 complex), we characterized the spectral profile as the mean frequency - instead of the maximum peak [20].

### Statistical Analyses

Correlations between SEEG and hd-EEG measures were performed with non-parametric Spearman’s correlation analyses. Differences among multiple groups were assessed with Kruskal-Wallis test (KW), followed by a post-hoc Wilcoxon Rank Sum test (WR, corrected for multiple comparisons using the False Discovery Rate method-FDR). Statistical interactions among stimulation parameters were performed with ANalysis Of VAriance (ANOVA). All descriptive values are reported along the manuscript as the mean ± standard deviation. All statistical analyses were performed in R.

## Results and Discussion

To the best of our knowledge, this is the first time that responses to intracortical SPES were studied simultaneously with SEEG and scalp hd-EEG, as previous concurrent recordings were only carried out for spontaneous activity while using low-density standard 10-20 systems [30–33].

Here, simultaneous scalp hd-EEG and intracranial SEEG recordings of CCEPs were performed in 36 awake drug-resistant epileptic patients undergoing SPES for presurgical evaluation (see Figure 1A-B-C). Overall, our dataset included 323 artefact-free recording sessions encompassing different stimulation parameters, which were clustered in three categories (see Figure 1D): *physical* (stimulation intensity and pulse width), *geometrical* (position of the bipolar contact with respect to grey/white matter and angle of the electrode with respect to the cortical surface), and *topological* (stimulated cortical area).

### General features of CCEPs were consistent between SEEG and hd-EEG

CCEPs were highly reproducible from trial to trial (Figure 1C) and characterized by a high signal-to-noise ratio both in intracerebral (SNR = 7.45 ± 10.73) and in hd-EEG (SNR = 6.96 ± 4.94) recordings (Figure 1D). We first quantified the overall strength of the response to SPES by computing GMFP (cumulated between 0ms and 500ms) at the hd-EEG level, and the percentage of significantly responding contacts at the SEEG level (Figure 2B). Then, we evaluated the waveshape of CCEPs by measuring the average amplitude of N1 and N2 across all sessions (Figure 2B and Methods). Despite displaying different waveshapes at the single contact level (see for example Figure 1E), hd-EEG and SEEG showed on average a similar waveform characterized by a prominence of the typical [27,34–36] N1 and N2 components (N1 = 9.5 ± 5.04μV; N2 = 11.36 ± 9.86μV for hd-EEG and N1 = 13.90 ± 7.04 z-value; N2 = 15.20 ± 7.02 z-value for SEEG; Figure 2A). Importantly, we found significant positive correlations (GMFP and number of responding contacts, *r*=0.592; N1 amplitude, *r*=0.313; N2 amplitude, *r*=0.553. All *p* <0.001) between the above-mentioned SEEG and hd-EEG measures (Figure 2C), suggesting that general features of CCEPs could be captured at both levels.

### Physical stimulation parameters: the effects of pulse intensity and width

The effects of varying stimulation intensity could be appreciated in the CCEPs when measuring N1, N2 and overall strength of the response both at the SEEG and hd-EEG level (Figure 3A). Statistical analysis performed with KW and post-hoc pairwise comparisons using WR tests revealed that the differences among the three stimulation intensities (≤1mA, 3mA and 5mA) could be fully captured by both SEEG and hd-EEG (Figure 3B). Specifically, SEEG showed a significant difference in the percentage of contacts responding to SPES (H_(2)_=25.70, *p*<0.001), in the N1 amplitude (H_(2)_=25.39, *p*<0.001) and in N2 amplitude (H_(2)_ =36.60, *p*<0.001). Similarly, hd-EEG showed significant differences in the GMFP (H_(2)_=10.05, *p*=0.006), in N1 amplitude (H_(2)_=12.13, *p*=0.002), as well as in N2 (H_(2)_=11.95, *p*=0.002). Post-hoc statistical analyses are reported in Table S2.

Conversely, differences in pulse width (0.5ms vs 1ms) were captured only by hd-EEG but not by SEEG (Figure 3C). Specifically, at the hd-EEG level, WR test showed that GMFP, amplitude of N1 and amplitude of N2 were significantly larger for 1ms than for 0.5ms pulse width (W=14193, W=13916, W=12663, respectively, all *p*<0.001; Figure 3D). Of note, both physical stimulation parameters (intensity and width) were not biased by any specific spatial distribution (see Figure S3A-B).

Overall, these results are in line with previous intracerebral studies which demonstrated that the amplitude of N1 and N2 components and, more in general, the amplitude of CCEPs depend on the amount of injected current [37-40]. However, while the effect of stimulation intensity has been clearly described, the effects of pulse width are less consistent across the literature [14,28,39,41]. Here, the larger hd-EEG responses elicited by longer pulse width stimulations may suggest the involvement of a larger network, implying broader polysynaptic activations [27] and recurrent activities [24,42]. In summary, complementing intracerebral explorations with whole brain hd-EEG measures confirms previous findings regarding stimulation intensity and suggests that the effects of pulse width may not be fully captured by SEEG recording alone.

### Geometrical stimulation parameters: the effects of contact position with respect to the cortex

First, we assessed the SEEG and hd-EEG responses to SPES when stimulating at different distances from grey-white matter interface (operationalized as GG/GW/WW; Figure 4A). We observed that this geometrical stimulation parameter affected CCEPs both at the SEEG and hd-EEG level, as assessed by KW statistical analyses. Specifically, this was true for all measures at the hd-EEG level (for GMFP, H_(2)_=15.03, *p*<0.001; for AMP N1, H_(2)_=26.41, *p*<0.001; for AMP N2 H_(2)_=11.95, *p*<0.01). Instead, at the SEEG level only the percentage of responding contacts and the amplitude of N2 showed a significant difference (H_(2)_=12.66, *p*<0.01; H_(2)_=17.47, *p*<0.001; respectively), while the N1 amplitude was not significantly affected (H_(2)_=5.12, *p*=0.077). In particular, except for N1 in SEEG, post-hoc comparisons showed that the stimulation of WW was more effective (*i.e*., larger CCEP response) with respect to the stimulation of GW, which in turn was more effective than the stimulation of GG (Figure 4B and Table S3).

**Figure 4.**
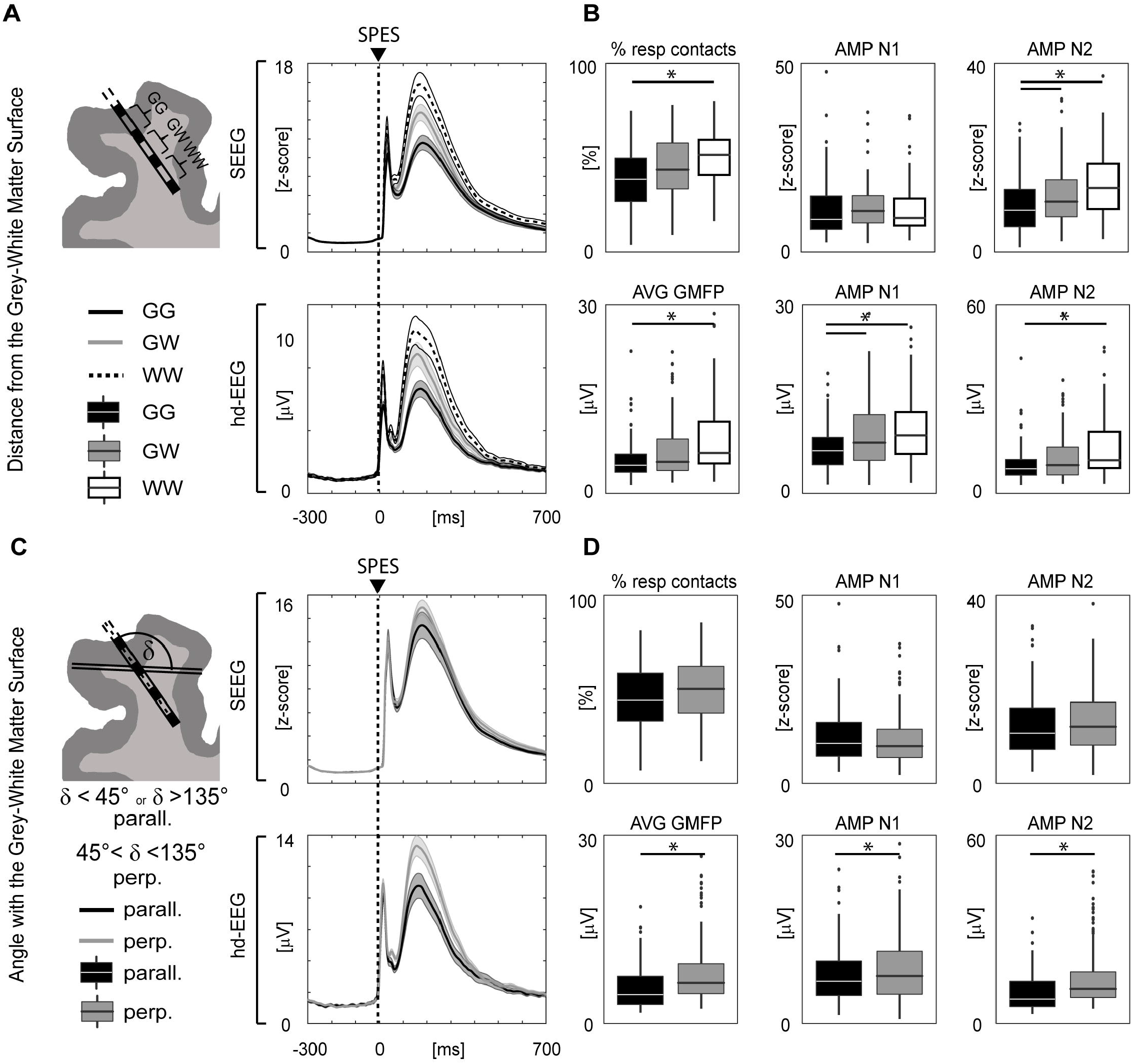
Hd-EEG and SEEG responses to SPES delivered at different geometrical stimulation parameters. **Panel A.** Left, outline of the distance from the gray-white matter surface (G and W respectively); right, grand average of data obtained from all the subjects and sessions. **Panel B**. *Upper line:* from left to right, percentage of responding contacts and N1 and N2 amplitudes for all sessions, recorded in SEEG. *Lower line:* from left to right, GMFP voltage and N1 and N2 amplitudes, for all sessions, recorded in hd-EEG. Asterisks indicate significant statistical differences (post-hoc, two-tailed WR, p<0.01, FDR corrected). **Panel C**. Left, outline of the angle with respect to the gray-white matter surface (parallel, perpendicular); right, grand average of data obtained from all the subjects and sessions. **Panel D.** *Upper line:* from left to right, percentage of responding contacts and N1 and N2 amplitudes for all sessions, recorded in SEEG. *Lower line:* from left to right, GMFP voltage and N1 and N2 amplitudes, for all sessions, recorded in hd-EEG. Asterisks indicate significant statistical differences (WR, p<0.01, FDR corrected).

The second geometrical parameter we considered was the angle with respect to the grey-white matter interface (operationalized as parallel/perpendicular, Figure 4C). In this case (Figure 3C), perpendicular stimulations led to significantly larger responses only at the hd-EEG level (for GMFP, W=6361 *p*=0.045; for AMP N1 W=6483 *p*=0.038; for AMP N2 W=6551 *p*=0.044). On the contrary, none of the SEEG measures showed significant differences (W=6588 *p*=0.125, W=5491 *p*=0.473, and W=5491 *p*=0.286 for percentage of responding contacts, AMP N1, and AMP N2, respectively). Of note, both geometrical parameters (white matter distance and angle) were not biased by any specific spatial distribution (Figure S3C-D).

Studies on intracerebral techniques that focused on the effect of geometrical stimulation parameters have been performed to optimize Deep Brain Stimulation protocols. According to these studies, small differences in electrode location [43–46], as well as orientation [47] can generate considerable differences in the activated white matter pathways. In line with these findings, the larger hd-EEG responses observed both with WW and perpendicular stimulations could be ascribed to the more extensive involvement of white-matter fiber bundles.

### Interactions between physical and geometrical stimulation parameters

The above-mentioned stimulation parameters could in principle interact at different levels. However, a model with the interaction of all the explored physical and geometrical parameters would require a larger sample. For this reason, we tested the interactions using pairwise bivariate ANOVAs. Overall, we observed significant interactions only at the hd-EEG level in pulse width/angle, pulse width/distance from white matter, intensity/distance from white matter (Figure S4). Specifically, the first two were found both for GMFP and amplitude of N2 while the latter was found only for N1 amplitude. Although it is conceivable that longer pulse widths and higher intensities might have stronger effects when delivered closer or perpendicular to white matter fiber bundles [48-51] and that this might reflect in more effective whole-brain level effects (*i.e*., recorded with hd-EEG), future studies including a larger sample size and a multivariate analysis would be needed to reach an exhaustive interpretation of these interactions.

### Topological stimulation parameters: the effect of stimulating different areas

Further, we evaluated whether the stimulation of different cortical areas was associated with differences in CCEP amplitude. At the hd-EEG level, we systematically observed that responses to the stimulation of the frontal cortex were larger than those obtained when stimulating any other cortical area. Specifically, as shown in Figure 5C, KW test and Wilcoxon Rank Sum post-hoc pairwise comparisons revealed a significant difference for all the measures at the hd-EEG level (GMFP: H_(5)_=15.45, *p*=0.008; AMP1 H_(5)_=20.32, *p*=0.001; AMP2 H_(5)_=19.85, *p*=0.001). On the contrary, among all the considered SEEG measures, only N1 showed a significant effect (percentage of responding contacts: H_(5)_=10.41, *p*=0.06; AMP1 H_(5)_=20.71, *p*=0.0009; AMP2 H_(5)_=9.29, *p*=0.09). Post-hoc statistical analyses are reported in Table S4.

**Figure 5.**
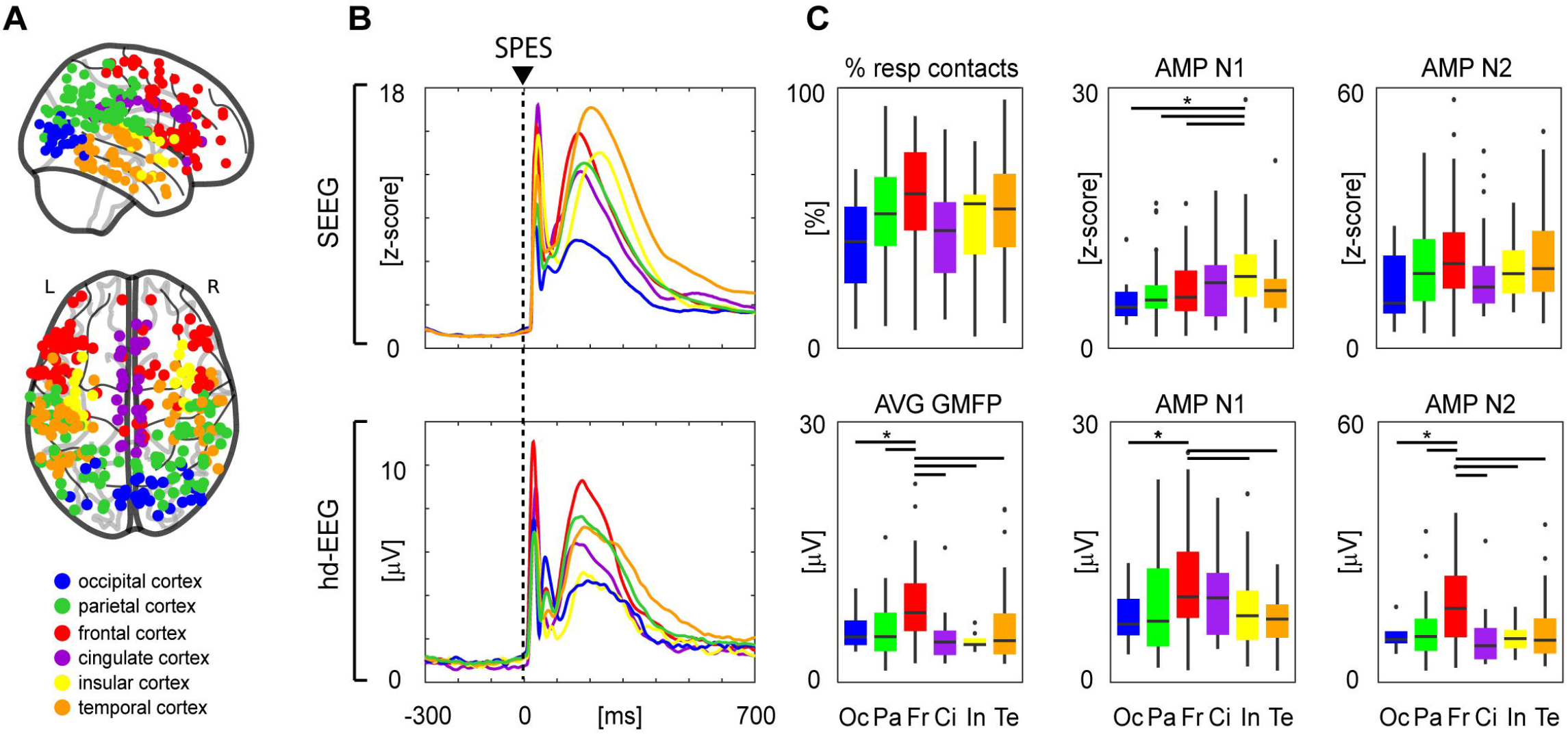
Hd-EEG and SEEG responses to SPES delivered through contacts in different cortical cortices (topological parameters). **Panel A.** Topographical distribution of the stimulations performed through bipolar SEEG contacts located in different cortical areas (cingulate cortex, insula, frontal cortex, occipital cortex, parietal cortex, temporal cortex). Color coding is consistent across the figure. **Panel B**. Grand average across sessions and subjects of the GMFP obtained by SPES of all the six cortices reported in Panel A; dashed vertical line indicates SPES timing. Here and across the figure, top panels refer to SEEG recordings, while bottom panels refer to hd-EEG recordings. **Panels C**. From left to right, boxplots of Average GMFP, amplitude of N1 and amplitude of N2. Asterisks indicate significant statistical differences obtained with a post-hoc, two-tailed WR test (p<0.05, FDR corrected).

High-amplitude responses to SPES of the frontal cortex could be due to the involvement of the circuits related to saliency [52,53], which are thought to be responsible for the generation of high amplitude scalp EEG graphoelements such as the K-complex [54] and the Vertex Wave [55]. Intriguingly, the latter is the largest graphoelement that can be evoked by sensory stimulation in an awake brain and is on average 25μV, with a peak-to-peak maximal amplitude of 35μV [52–55].

### Comparing invasive and non-invasive brain stimulation techniques

In our dataset, CCEPs voltage at Cz were on average 43.42μV (average reference; 53.1μV when referenced to mastoid), reaching a peak-to-peak maximum of 172.16μV (average reference; 214.43μV when referenced to mastoid), and thus much larger than any sensory evoked potential recorded during wakefulness. This finding is particularly interesting when considering CCEPs impact on the subjects’ awareness: despite eliciting massive and long-lasting activations of cortical circuits, none of our intracranial stimulation resulted in any reportable perceptual event.

Interestingly, CCEPs’ voltages at the scalp EEG level were also out of scale with respect to the scalp EEG responses typically obtained when perturbing similar circuits with non-invasive peripheral or direct stimulation in awake healthy subjects. Indeed, sensory (be it auditory, somatosensory or visual) evoked potentials may range from a fraction of a microvolt to few microvolts, while TMS evoked potentials (TEPs) may reach amplitudes of about 20μV [56-59]. Importantly, our results showed that, similarly to TMS, SPES could elicit large EEG components that persist for hundreds of milliseconds, thus corroborating the idea that late components genuinely reflect the effects of direct cortical rather than peripheral activation [60].

Furthermore, combining hd-EEG with SPES allowed to directly compare invasive and non-invasive (TMS) stimulation methods in terms of spectral properties emerging at the local and at whole brain level when perturbing the brain at different sites - as in TMS-EEG investigations [20]. To this aim, we calculated time-frequency spectra (ERSP) of the CCEPs collected with SEEG and hd-EEG when stimulating occipital, parietal or frontal cortices (Figure 6A-B). Then, we cumulated the ERSPs over time (between 5ms and 150ms) to obtain a spectral profile for each session, whose grand average is depicted in Figure 6C-D. This analysis showed that the CCEPs evoked by the stimulation of the occipital, parietal and frontal cortices were characterized by a rostro-caudal gradient of mean frequencies - *i.e*., occipital<parietal<frontal (Figure 6C-D). These differences were significant both at the hd-EEG and at the SEEG level as assessed by KW tests (hd-EEG: H_(2)_=16.49, p=0.0002; SEEG: H_(2)_=8.31, p=0.013) and post-hoc one-tailed WR tests (Table S5). These results obtained with intracortical SPES confirm previous non-invasive TMS-EEG assessments [20,56] and reflect site specific spectral properties that are amplitude-independent and can be observed both at the whole brain level (*i.e*., with the scalp EEG) as well as locally (SEEG).

**Figure 6.**
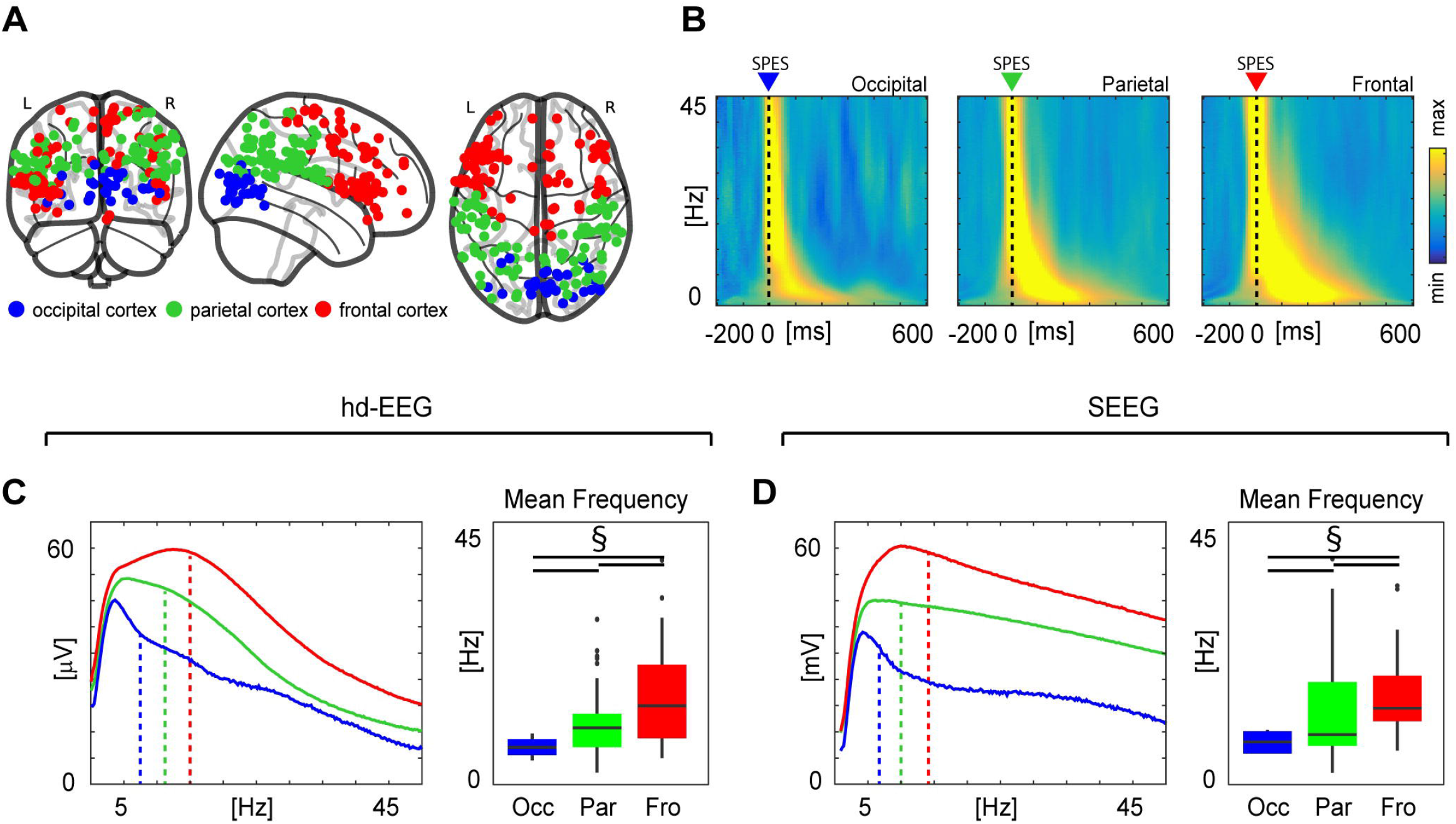
Reproducing TMS-EEG experiments: rostro-caudal gradient of cortical spectral features. **Panel A.** Topographical distribution of the stimulations performed through bipolar SEEG contacts located in different cortices (occipital cortex, parietal cortex, frontal cortex). Color coding is consistent across the figure. **Panel B.** The spectral properties (ERSP) emerged at the whole brain level after SPES in the three different sites (occipital, parietal, frontal), recorded with hd-EEG. **Panels C** concerns hd-EEG recordings. *Left*. Grand average across sessions and subjects of spectral profile, namely ERSPs cumulated over time (between 5ms and 150ms), obtained by SPES of occipital, parietal and frontal cortices; dashed vertical lines indicate mean frequencies. *Right*. Boxplot of the mean frequency for occipital, parietal, and frontal cortices. § indicates significant statistical differences (p<0.05) obtained with one-tailed Wilcoxon test, under the assumption that the mean frequency of occipital, parietal and frontal cortex are characterized by a rostro-caudal gradient [22]. **Panel D.** Same as C but concerning SEEG recordings.

### Limitations

Our results were obtained from a population of epileptic patients whose clinical condition and specific treatment [61] may affect both invasive and non-invasive recordings. To minimize this confound, we did not include any SEEG contact located in the SOZ (as verified by surgical resection) or exhibiting sustained pathological interictal activity. Moreover, we excluded from the analyses all the CCEPs showing evoked epileptic activity at the SEEG and/or at the hd-EEG level [9].

Clinical needs also constrained the exploration of physical stimulation parameters to few pulse intensities and two pulse widths. Future studies encompassing multiple intensity and pulse width steps, like Paulk and colleagues [63], will allow for a more systematic comparison between invasive and non-invasive stimulation techniques.

Finally, the combination of SEEG and hd-EEG entails specific data acquisition protocols to prevent infective risks. This implies a short duration of SPES procedures and thus the acquisition of few trials from a limited number of sites per patient. To compensate for these constraints, we verified that the number of acquired trials led to reliable responses in terms of SNR - both at the SEEG and hd-EEG level - and we included a relatively large number of stimulation sessions (N=323) and subjects (N=36). This sample size allowed to perform univariate analyses and to assess interactions through bivariate models (Figure S4). Larger datasets will ensure the possibility of performing multivariate analyses considering all the explored variables. In this respect, the results shown in the present manuscript represent a first step toward a more comprehensive description of the scalp EEG responses to SPES and their relationship with intracerebral recordings.

### Conclusions

The present results show that CCEPs recorded with hd-EEG are overall aligned with those obtained with invasive SEEG recordings. Most important, they show that macroscale hd-EEG recordings are exquisitely sensitive to variations in stimulation parameters, including local changes in physical and geometrical stimulus properties, while overcoming the limitations typical of sparse recordings.

In general, the possibility of studying and comparing across subjects the effects of multiple local intracortical perturbations at the whole brain level opens interesting fields of investigations. For example, it may complement current datasets on the structural [64] and functional [65] connectomes with an effective connectome [28] whereby intracortical interactions are systematically studied by a causal perspective in the common recording space of scalp EEG and with a full assessment of spatio-temporal dynamics. Moreover, hd-EEG recordings allow direct comparisons between the CCEPs and classic evoked potentials elicited by non-invasive stimulation. For example, EEG responses to SPES reproduced the rostro-caudal spectral gradient as previously shown by TMS-EEG measurements and were found to be systematically larger than any sensory evoked potential that can be elicited during wakefulness, including those associated with stimulus perception. Along these lines, future studies should also explore, in terms of their whole-brain spatio-temporal features, why some brain responses are associated with perceptual events and others do not.

## Authors contributions

Conceptualization: S.P., E.M., S.R. and A.P.; supervision: A.P. and M.M.; data collection: S.P., S.R. A.P., F.Z., A.R., and I.S. data curation: E.M. and AP; data analyses: S.P., E.M., A.P., S.R., M.F., A.C.; clinical investigation: I.S., J.L., D.G., P.dO, statistical analysis: E.M., S.P., S.R. and A.P.; writing original draft: S.P., A.P., E.M., S.R, and S.S.; review and editing: all authors.

## Declarations of interest

None of the authors have conflicts of interest to disclose in relationship with the current work.

## Funding

This research has received funding from the European Union’s Horizon 2020 Framework Programme for Research and Innovation under the Specific Grant Agreement No. 720270 (Human Brain Project SGA1-to M.M.), No. 785907 (Human Brain Project SGA2 - to M.M. and P.A.), and No. 945539 (Human Brain Project SGA3 - to M.M. and P.A.). The study has also been partially funded by the grant “Sinergia” CRSII3_160803/1 of the Swiss National Science Foundation (to M.M.) and the Tiny Blue Dot Foundation (to M.M.).

## Acknowledgements

We are grateful to Mario Rosanova, Susanna Bianchi, Jacopo Favaro, Michela Solbiati, Aurora Sium, Alessandro Viganò, Renzo Comolatti, Giulia Furregoni and Martina Revay for their comments and suggestions.

## Notes

### Competing Interest Statement

The authors have declared no competing interest.

