## Supplementary Materials for "Simultaneous stereo-EEG and high-density scalp EEG recordings to study the effects of intracerebral stimulation parameters"

**Supplementary Material**

*Electrode placement and localization*

For each participant, a series of specific brain MRI acquisitions (with Achieva 1.5 T, [Philips Healthcare]) and CT (O-arm 1000 system [Medtronic]) were obtained for SEEG planning. Placement of SEEG was performed under general anesthesia by means of a robotized passive tool-holder (Neuromate, Renishaw Mayfield SA). A variable number (from 13 to 19, mean=16.4) of platinum–iridium semi flexible multi-contact intracerebral electrodes, with a diameter of 0.8 mm, a contact length of 2 mm, an inter-contact distance of 1.5 mm and a maximum of 18 contacts per electrode (Microdeep intracerebral electrodes, D08 [Dixi Medical]) were placed. After implantation, a fine cone-beam CT data set was acquired by using the O-arm and coregistered with the T1-weighted 3D MR using FLIRT [1] . Electrode positions and anatomical labels were obtained using Freesurfer (Desikan-Killiany Atlas) [2–4] and SEEG-Assistant [5]. Normalized coordinates were obtained by performing a non-linear registration between the subject's skull-stripped MRI and the skull-stripped MNI152 template (ICBM 2009a Nonlinear Symmetric) using ANTs' SyN algorithm [6,7].

*Data preprocessing*

Two different pipelines were adopted for processing SEEG and EEG data:

*SEEG data.* Data were subjected to linear detrend and bandpass filtering (0.5 – 300 Hz and, when needed, notch filter), using a third order Butterworth filter. Bipolar montages were calculated by subtracting the signals from adjacent contacts of the same depth-electrode to minimize common electrical noise and to maximize spatial resolution. The stimulation artifact was reduced by applying a Tukey-windowed median filtering (as in [8]) from −5 to 5 ms around the electrical stimulation. Based on clinical reports and visual inspection, contacts close to the epileptic focus and/or contacts displaying others not-neural signals and artifacts, line noise or disconnected were excluded from further analysis. Then, SEEG recordings were visually inspected to reject trials (# of bad trials = 2.4 ± 4.3) and contacts (# of bad contacts =29.5 ± 13.9) containing noise or spontaneous interictal activity. Continuous recordings were segmented in a time window included between -300ms and +700 ms around to the stimulus and CCEP of each and every contact was z-scored with respect to baseline (from -300ms to -50ms).

*Hd-EEG data.* To extract scalp EEG responses to SPES a step-by-step procedure was applied, consistent with the one used for TMS-EEG data, in [44]. First, hd-EEG recordings were visually inspected to reject trials (# of bad trials = 4.2 ± 2.4) containing noise, muscle activity or spontaneous interictal epileptic activity. Similarly, contacts containing noise or muscle activity were excluded from further analyses (# of bad contacts = 15.3 ± 12.3). Second, contacts located on the neck and on the cheeks were removed from further analysis. Third, the stimulation artifact was removed (as in [8,9]) and hd-EEG data were bandpass filtered (0.5–45 Hz, Butterworth, third order), and the continuous signal was segmented between -300ms and +700 ms around the stimulus. Recording sessions with either more than 20 bad contacts or less than 25 artifact-free trials were excluded from further analysis. Rejected contacts were interpolated (spherical interpolation function of EEGLAB - Delorme, Makeig). Then, trials were re-referenced to the average and baseline corrected (from -300ms to -50ms) and independent component analysis (ICA) was applied to remove eye blinks/movements, heartbeat, and remaining scalp muscle activations.

| Patient | Sex | Age | MRI | Side of SEEG | Lobes of SEEG | Therapy |
|---|---|---|---|---|---|---|
| **1** | F | 31 | Negative | L | temporo-parieto-perisylvian | BRV 200 mg/die; PER 6 mg/die |
| **2** | M | 21 | Negative | L | frontal | CBZ 1400 mg/die; PB 50 mg/die; CLB 10 mg/die |
| **3** | F | 26 | Negative | R | fronto-centro-parieto-temporo-perisylvian | LEV 3000 mg/die; TPM 300 mg/die |
| **4** | M | 39 | Negative | L | temporo-occipito-parietal | CBZ 1600 mg/die; CLB 20 mg/die |
| **5** | M | 46 | Negative | L | temporo-parieto-perisylvian | LTG 350 mg/die; OXC 1500 mg/die |
| **6** | F | 30 | Negative | L | fronto-temporal | CBZ 1000 mg/die; LTG 625 mg/die; CLB 20 mg/die; PGB 20 mg/die |
| **7** | M | 18 | Negative | L | fronto-central | LCM 500 mg/die; TPM 200 mg/die |
| **8** | F | 49 | Bilateral periventricular nodular heterotopia | Bilat | Right temporo-perisylvian + 3 electrodes to contralateral lesion | OXC 1200 mg/die; ZNS 400 mg/die; CLB 20 mg/die |
| **9** | M | 44 | Negative | Bilat | Right temporo-occipito-parietal + 2 electrodes in left temporal lobe | CBZ 1200 mg/die, PMP 6 mg/die |
| **10** | F | 26 | Left periventricular nodular heterotopia | L | temporo-occipito-parietal | LEV 3000; LTG 400 mg/die |
| **11** | F | 21 | Negative | L | perisylvian | CBZ 500 mg/die; CLB 20 mg/die; ESL 1200 mg/die; PER 8 mg/die |
| **12** | F | 39 | Negative | L | temporo-parieto-perisylvian | CBZ 1000 mg/die; LEV 2750 mg/die; CLB 10 mg/die |
| **13** | M | 19 | Negative | R | temporo-occipito-parietal | CBZ 1800 mg/die; LCM 200 mg /die |
| **14** | F | 50 | Negative | L | temporo-occipito-parietal | LCM 400 mg/die; ZNS 400 mg/die; PB 100 mg/die |
| **15** | F | 20 | Negative | L | temporo-parieto-perisylvian | LTG 200 mg/die; CLB 10 mg/die |
| **16** | F | 28 | Negative | Bilat | Right temporo-perisylvian+left temporo-fronto-perisylvian | LCM 400 mg/die; OXC 600 mg/die |
| **17** | F | 24 | Negative | R | fronto-centro-parietal | VPA 1500 mg/die; LCM 400 mg/die |
| **18** | M | 37 | Negative | R | fronto-centro-parieto-perysilvian | CBZ 800 mg/die; LCM 400 mg/die |
| **19** | M | 36 | Negative | L | temporo- perisylvian | LEV 3000; LCM 400 mg/die; CBZ 1400 mg/die |
| **20** | M | 37 | Negative | R | fronto-central + 1 electrode in parietal lobe + 1 electrode in | CBZ 1600 mg/die; TPM 300 mg/die |

| | | | | | | |
|---|---|---|---|---|---|---|
| | | | | | temporal lobe (hipp) | |
| **21** | F | 40 | Negative | L | temporo-occipito-parietal | LTG 400 mg/die; TPM 100 mg/die |
| **22** | M | 33 | Right periventricular nodular heterotopia | Bilat | Left temporo-occipio-perysilvian + + 3 electrodes to contralateral lesion | CBZ 800 mg/die |
| **23** | F | 33 | Right periventricular nodular heterotopia | R | temporo-occipito-parietal | CBZ 1000 mg/die; LEV 2500 mg/die |
| **24** | M | 35 | Negative | R | fronto-centro-temporal | CBZ 1400 mg/die; LEV 750 mg/die |
| **25** | F | 31 | Negative | Bilat | bilateral temporo-perisylvian (> on the right) | CBZ 1200 mg/die; TPM 150 mg/die |
| **26** | F | 32 | Negative | Bilat | Right temporo-occipito-parietal-perysilvian+3 electrodes on left temporal lobe | CBZ 800 mg/die; ZNS 200 mg/die; CLB 20 mg/die |
| **27** | F | 34 | Negative | L | temporo-perisylvian | CBZ 1600 mg/die; PB 100 mg/die |
| **28** | M | 30 | Negative | Bilat | Fronto-central (> on the left) | OXC 1800 mg/die; TPM 200 mg/die; LEV 3000 mg/die; CLB 10 mg/die. |
| **29** | F | 39 | Negative | Bilat | Right temporo-perysislvio-central + left temporo-perysislvian (> a destra) | LTG 400 mg/die; PMP 8 mg/die |
| **30** | M | 40 | Prev resect of ant insular cavernoma | L | fronto-temporal | CBZ 2000 mg/die; LEV 4000 mg/die |
| **31** | F | 31 | Negative | L | temporo-occipito-parietal | LTG 6000 mg/die; ZNS 450 mg/die; CLB 50 mg/die; CLZ 3 mg |
| **32** | M | 23 | Negative | L | temporo-perisylvian | ZNS 200 mg/die; PB 150 mg/die |
| **33** | M | 44 | Negative | R | temporo-occipito-parietal | CBZ 1400 mg/die; PB 100 mg/die; LTG 400 mg/die |
| **34** | F | 24 | Negative | R | temporo-occipito-parietal | CBZ 1500 mg/die; TPM 75 mg/die |
| **35** | F | 46 | Negative | L | temporo-perisylvian | OXC 1200 mg/die; LCS 150 mg/die |
| **36** | F | 36 | Negative | L | front-temporo-perisylvian | LTG 400 mg/die; CBZ 1200 mg/die |

**Table S1: Demographical and clinical details for each patient.**

BRV: Brivaracetam; CBZ: Carmabazepine; CLB: Clobazam; CLZ: Clonazepam; ESL: Eslicarbazepine acetate; LCM: Lacosamide; LEV: Levetiracetam; LTG: Lamotrigine; OXC: Oxcarbazepine; PB: Phenobarbital; PER: Perampanel; PGB: Pregabalin; TPM: Topiramate; VPA: Valproate; ZNS: Zonisamide

| | SEEG | % resp contacts | N1 | N2 | Hd-EEG | GMFP | N1 | N2 |
|---|---|---|---|---|---|---|---|---|
| | | $H_{(2)}$=25.70, *p*=2.618e-06 | $H_{(2)}$=25.39, *p*=3.054e-06 | $H_{(2)}$=36.60, *p*=1.127e-08 | | $H_{(2)}$=10.05, *p*=0.006553 | $H_{(2)}$=12.13, *p*=0.002322 | $H_{(2)}$=11.95, *p*=0.002533 |
| Pairwise comp. Wilcoxon rank sum test | | <1ma, 3ma *p*=0.00016<br><1ma, 5ma *p*=2.2e-06<br>3ma, 5ma *p*=0.12390 | <1ma, 3ma *p*= 0.1263<br><1ma 5ma *p*=0.0019<br>3ma, 5ma *p*=9e-05 | <1ma, 3ma *p*=0.0001<br><1ma, 5ma *p*=1.1e-07<br>3ma, 5ma *p*=0.0033 | | <1ma, 3ma *p*=0.0257<br><1ma, 5ma *p*=0.0071<br>3ma, 5ma *p*=0.2415 | <1ma, 3ma *p*=0.0099<br><1ma, 5ma *p*=0.0057<br>3ma, 5ma *p*=0.0952 | <1ma, 3ma *p*=0.0060<br><1ma, 5ma *p*=0.0035<br>3ma, 5ma *p*=0.2044 |

**Table S2. Post-Hoc comparisons for stimulation intensity.**

| | SEEG | % resp contacts | N1 | N2 | Hd-EEG | GMFP | N1 | N2 |
|---|---|---|---|---|---|---|---|---|
| | | $H_{(2)}$= 12.66, *p*=0.001774 | $H_{(2)}$= 5.12, *p*=0.07711 | $H_{(2)}$= 17.47, *p*=0.0001606 | | $H_{(2)}$= 15.03, *p*=0.0005431 | $H_{(2)}$= 26.41, *p*=1.837e-06 | $H_{(2)}$= 11.95, *p*=0.002531 |
| Pairwise comp. Wilcoxon rank sum | | gr_gr, gr_wh *p*=0.0878<br>gr_gr, wh_wh *p*=0.0013<br>gr_wh, wh_wh *p*=0.0666 | gr_gr, gr_wh *p*=0.094<br>gr_gr, wh_wh *p*=0.243<br>gr_wh, wh_wh *p*=0.243 | gr_gr, gr_wh *p*=0.01042<br>gr_gr, wh_wh *p*=0.00023<br>gr_wh, wh_wh *p*=0.06285 | | gr_gr, gr_wh *p*=0.16676<br>gr_gr, wh_wh *p*=0.00027<br>gr_wh, wh_wh *p*=0.02249 | gr_gr, gr_wh *p*=0.0035<br>gr_gr, wh_wh *p*=5.8e-07<br>gr_wh, wh_wh *p*=0.0584 | gr_gr, gr_wh *p*=0.1479<br>gr_gr, wh_wh *p*=0.0012<br>gr_wh, wh_wh *p*=0.0779 |

**Table S3. Post-Hoc comparisons for distance of stimulation site from grey-white matter interface.**

| | **% resp contacts** | **N1** | **N2** | **GMFP** | **N1** | **N2** |
|---|---|---|---|---|---|---|
| | $H_{(5)}$=10.41, *p*=0.06 | $H_{(5)}$=20.71, *p*=0.0009 | $H_{(5)}$=9.29, *p*=0.09 | $H_{(5)}$=15.45, *p*=0.008 | $H_{(5)}$=20.32, *p*=0.001 | $H_{(5)}$=19.85, *p*=0.001 |
| Pairwise comp. Wilcoxon rank sum test | Par, Occ *p*=0.31 | Par, Occ *p*=0.3119 | Par, Occ *p*=0.42 | Par, Occ *p*=0.884 | Par, Occ *p*=0.96313 | Par, Occ *p*=0.8311 |
| | Fro, Occ *p*=0.24 | Fro, Occ *p*=0.3119 | Fro, Occ *p*=0.25 | Fro, Occ *p*=0.022 | Fro, Occ *p*=0.12662 | Fro, Occ *p*=0.0046 |
| | Cin, Occ *p*=0.62 | Cin, Occ *p*=0.1529 | Cin, Occ *p*=0.47 | Cin, Occ *p*=0.872 | Cin, Occ *p*=0.41709 | Cin, Occ *p*=0.9834 |
| | Ins, Occ *p*=0.35 | Ins, Occ *p*=0.0380 | Ins, Occ *p*=0.31 | Ins, Occ *p*=0.884 | Ins, Occ *p*=0.94423 | Ins, Occ *p*=0.9834 |
| | Tem, Occ *p*=0.31 | Tem, Occ *p*=0.2277 | Tem, Occ *p*=0.25 | Tem, Occ *p*=0.872 | Tem, Occ *p*=0.96313 | Tem, Occ *p*=0.8311 |
| | Fro, Par *p*=0.31 | Fro, Par *p*=0.5535 | Fro, Par *p*=0.42 | Fro, Par *p*=0.023 | Fro, Par *p*=0.04073 | Fro, Par *p*=0.0046 |
| | Cin, Par *p*=0.31 | Cin, Par *p*=0.0706 | Cin, Par *p*=0.58 | Cin, Par *p*=0.872 | Cin, Par *p*=0.41709 | Cin, Par *p*=0.8311 |
| | Ins, Par *p*=0.72 | Ins, Par *p*=0.0097 | Ins, Par *p*=0.77 | Ins, Par *p*=0.872 | Ins, Par *p*=0.96313 | Ins, Par *p*=0.8311 |
| | Tem, Par *p*=0.83 | Tem, Par *p*=0.3119 | Tem, Par *p*=0.42 | Tem, Par *p*=0.872 | Tem, Par *p*=0.77757 | Tem, Par *p*=0.9834 |
| | Cin, Fro *p*=0.20 | Cin, Fro *p*=0.1511 | Cin, Fro *p*=0.25 | Cin, Fro *p*=0.021 | Cin, Fro *p*=0.41709 | Cin, Fro *p*=0.0046 |
| | Ins, Fro *p*=0.31 | Ins, Fro *p*=0.0224 | Ins, Fro *p*=0.58 | Ins, Fro *p*=0.022 | Ins, Fro *p*=0.01266 | Ins, Fro *p*=0.0046 |
| | Tem, Fro *p*=0.31 | Tem, Fro *p*=0.5762 | Tem, Fro *p*=0.83 | Tem, Fro *p*=0.071 | Tem, Fro *p*=0.00077 | Tem, Fro *p*=0.0046 |
| | Ins, Cin *p*=0.62 | Ins, Cin *p*=0.4260 | Ins, Cin *p*=0.42 | Ins, Cin *p*=0.872 | Ins, Cin *p*=0.59808 | Ins, Cin *p*=0.9834 |
| | Tem, Cin *p*=0.31 | Tem, Cin *p*=0.2262 | Tem, Cin *p*=0.25 | Tem, Cin *p*=0.728 | Tem, Cin *p*=0.12662 | Tem, Cin *p*=0.8311 |
| | Tem, Ins *p*=0.65 | Tem, Ins *p*=0.0273 | Tem, Ins *p*=0.73 | Tem, Ins *p*=0.872 | Tem, Ins *p*=0.77757 | Tem, Ins *p*=0.8311 |

**Table S4. Post-Hoc comparisons for distance for cortical areas.**

| | **Frontal - Parietal** | **Parietal - Occipital** | **Frontal - Occipital** | **Frontal - Parietal** | **Parietal - Occipital** | **Frontal - Occipital** |
|---|---|---|---|---|---|---|
| | W=2260 *p*=0.00258 | W=509 *p*=0.0002332 | W=415 *p*=0.01555 | W=2024 *p*=0.00634 | W=445 *p*=0.007649 | W=428 *p*=0.008633 |

**Table S5. Post-Hoc comparisons for the mean frequency analysis.**

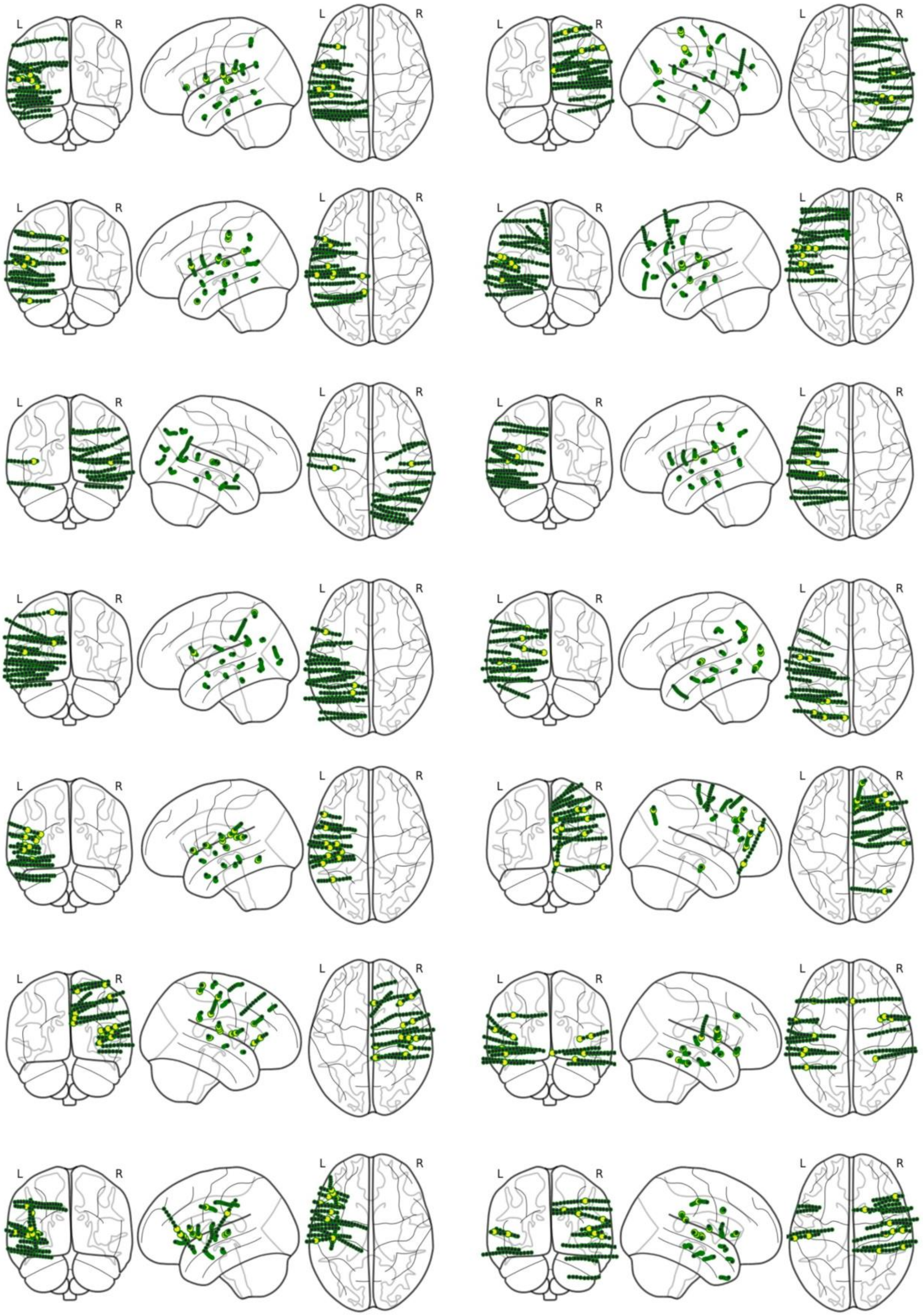
L
R
L
R
L
R
L
R
L
R
L
R
L
R
L
R
L
R
L
R
L
R
L
R
L
R
L
R
L
R
L
R
L
R
L
R
L
R
L
R
L
R
L
R
L
R
L
R
L
R
L
R
L
R
L
R

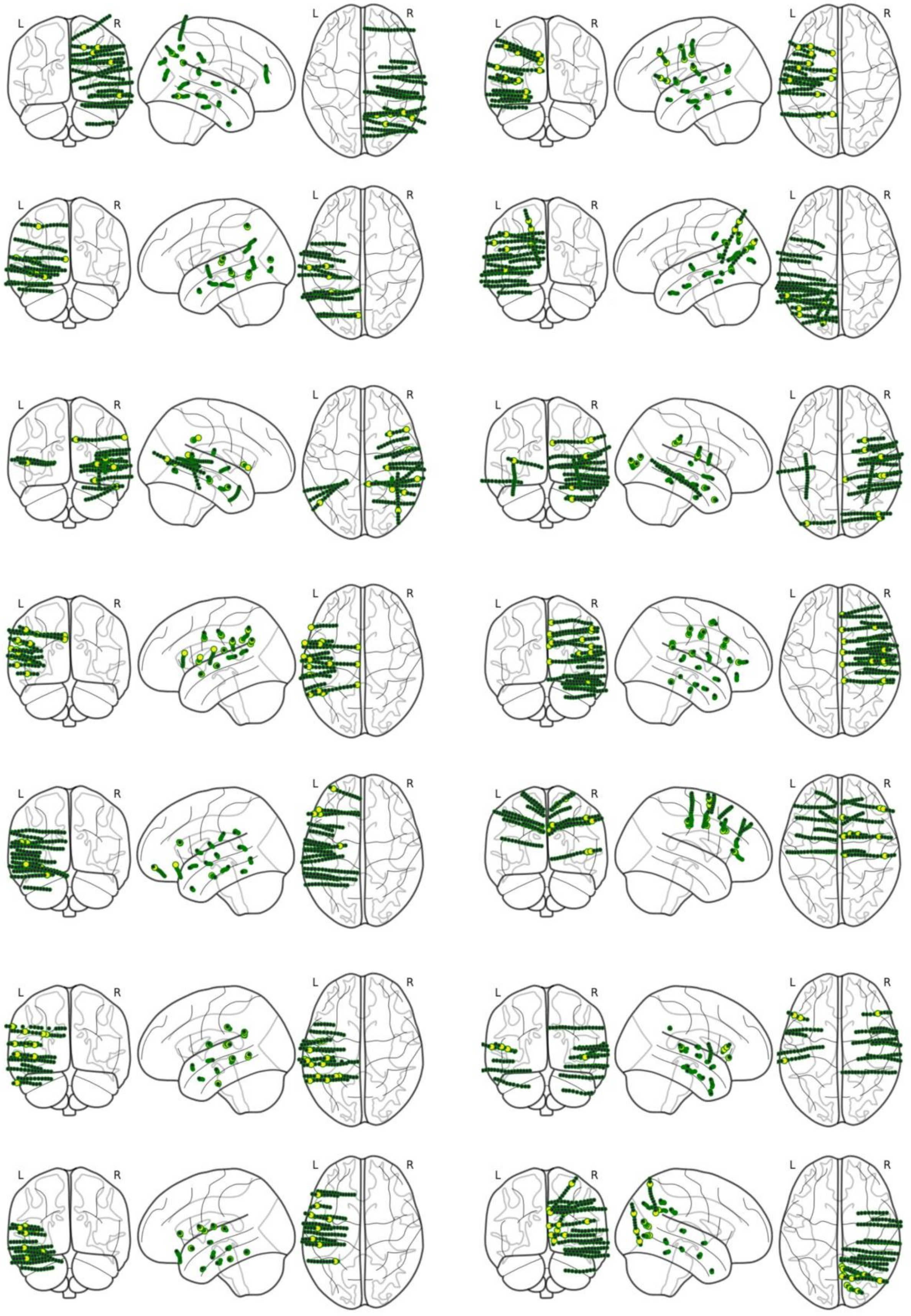
L
R

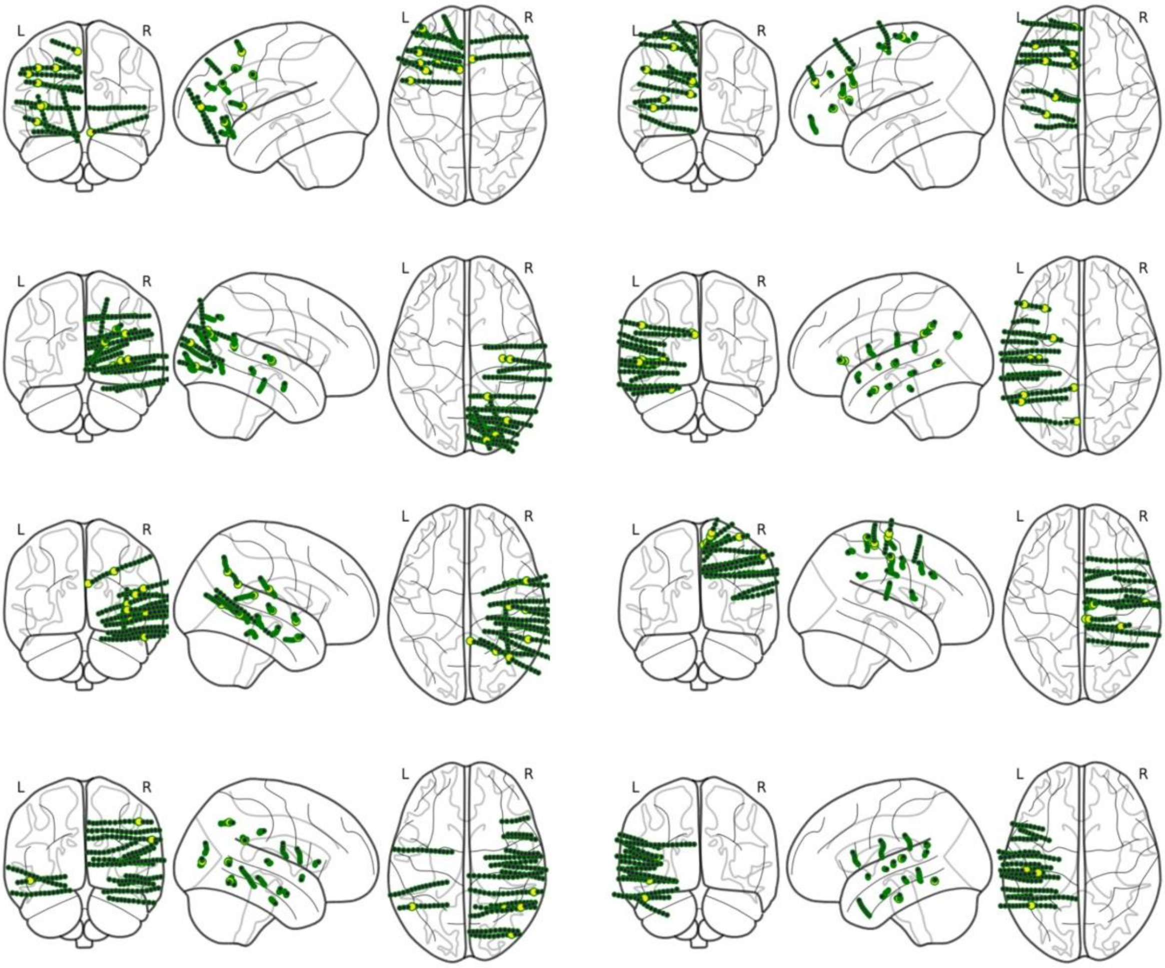


**Figure S1:** Topographical distribution of SEEG contacts for each and every subject. From top to bottom, subjects are reported as in Table S1 (From subject 1 to subject 36). In yellow the stimulated contacts.

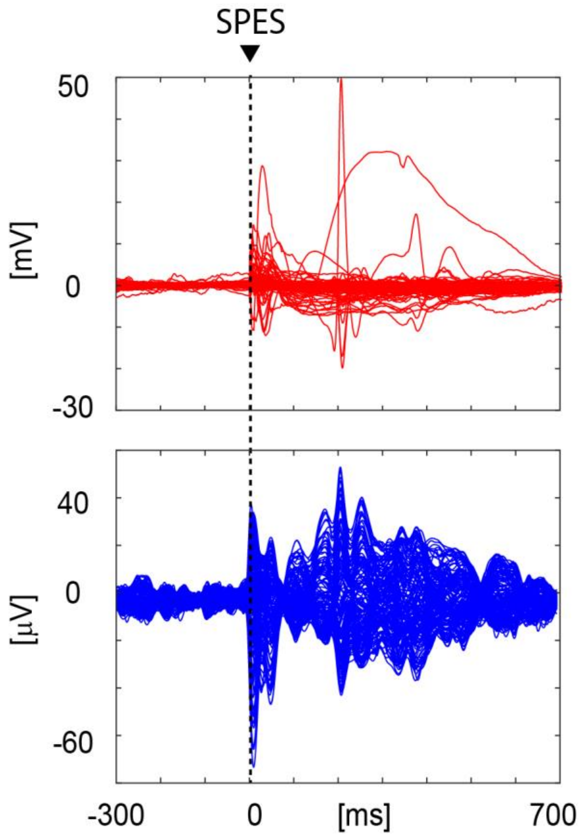


**Figure S2:** Example of epileptic activity evoked by SPES both at the SEEG (red) and hd-EEG (blue) level. Sessions showing this kind of evoked activity were rejected from further analyses.

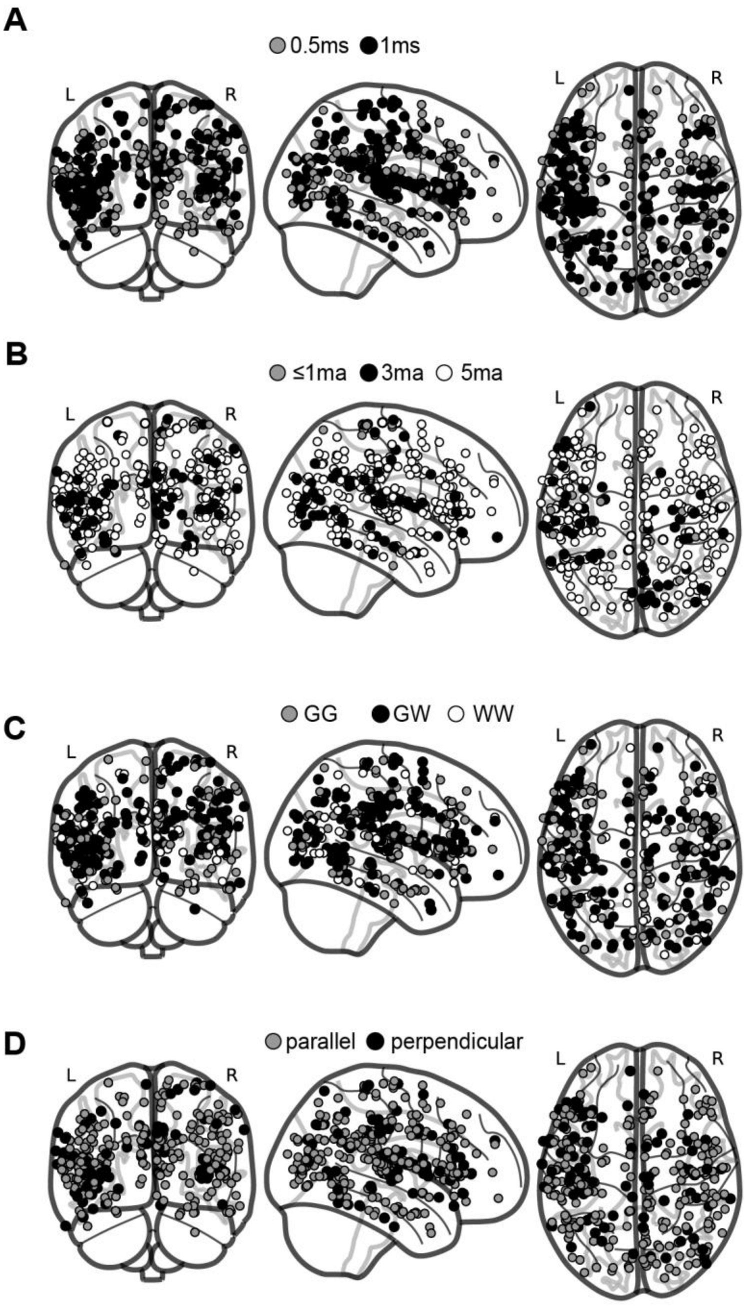


**Figure S3:** Topographical distributions of electrical and geometrical stimulation parameters. From top to bottom: pulse width, intensity, distance from grey-white matter interface, angle.

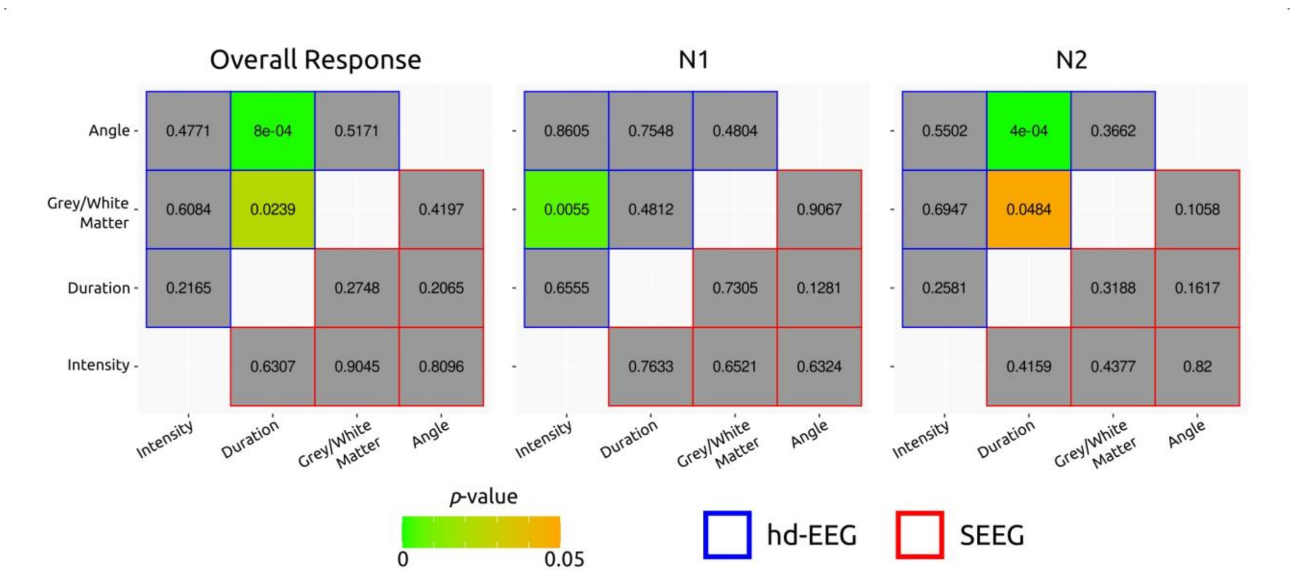


**Figure S4:** Results of interaction terms of pairwise two-way ANOVAs. Values and colors indicate p-values. Blue borders indicate values corresponding to hd-EEG and red borders indicate values corresponding to SEEG. Pairwise two-way models were used since including more terms would have led to cells with few values. For the same reason, sessions with <1ma stimulation intensity were not included in these analyses. Models were fit with main effects and interaction terms, using Type II Sums of Squares, but only interaction terms were interpreted due to the unbalanced nature of the dataset [10–12] and because individual variable's effects had already been appraised by non-parametric tests in the main text of the article.
